# Kaiju: Fast and sensitive taxonomic classification for metagenomics

**DOI:** 10.1101/031229

**Authors:** Peter Menzel, Kim Lee Ng, Anders Krogh

## Abstract

The constantly decreasing cost and increasing output of current sequencing technologies enable large scale metagenomic studies of microbial communities from diverse habitats. Therefore, fast and accurate methods for taxonomic classification are needed, which can operate on increasingly larger datasets and reference databases. Recently, several fast metagenomic classifiers have been developed, which are based on comparison of genomic *k*-mers. However, nucleotide comparison using a fixed *k*-mer length often lacks the sensitivity to overcome the evolutionary distance between sampled species and genomes in the reference database. Here, we present the novel metagenome classifier `Kaiju` for fast assignment of reads to taxa. Kaiju finds maximum exact matches on the protein-level using the Borrows-Wheeler transform, and can optionally allow amino acid substitutions in the search using a greedy heuristic. We show in a genome exclusion study that Kaiju can classify more reads with higher sensitivity and similar precision compared to fast *k*-mer based classifiers, especially in genera that are underrepresented in reference databases. We also demonstrate that Kaiju classifies more than twice as many reads in ten real metagenomes compared to programs based on genomic *k*-mers. Kaiju can process up to millions of reads per minute, and its memory footprint is below 6 GB of RAM, allowing the analysis on a standard PC. The program is available under the GPL3 license at: http://bioinformatics-centre.github.io/kaiju

## Introduction

Using random DNA shotgun sequencing, it is possible to directly obtain total genomic DNA from an environmental sample without the need for laboratory cultures. This “metagenomic” approach has become a standard method for characterizing the biodiversity, gene contents, and metabolic processes of bacterial and archaeal communities and is used on increasingly larger scales [20, 22, 21]. Due to decreasing costs of high-throughput sequencing and the recent revelations of the importance of microbiomes for health and disease [12, 24], metagenomic analyses are also likely to become part of routine clinical diagnostics and detection of pathogens.

One of the major biological questions in metagenomics is the inference of the composition of a microbial community, *i.e.,* the relative abundances of the sampled organisms. Hence, given random shotgun sequencing reads, the underlying computational problem is the assignment of individual reads to taxa, usually by comparison to a reference database. Traditionally, this task is solved by local sequence alignment either on nucleotide level, when comparing sequencing reads to a database of microbial genomes, or on protein level when translating reads to amino acid sequences and comparing to a catalog of microbial genes. However, with increasing volumes of microbial genome databases and se output, computational methods need to catch up, as traditional methods based on local sequence alignment are too slow in order to cope with the increasing amount of data.

For the similar problem of mapping sequencing reads to a reference genome, heuristic methods achieve speed improvements of orders of magnitude by using advanced index structures for fast identification and extension of short exact matches (seeds) between query and a reference genome [7]. However, these mappers are not suited for classification of metagenomic sequences, because they only work on DNA in a usually semi-global alignment model and assume near-identity of read sequences and reference genome,

Thus, programs have been developed for fast taxonomic classification of individual sequencing reads by using hash-based index structures built from a set of reference sequences, typically a database of complete microbial genomes. To achieve a high speed, these algorithms do not use traditional local alignment methods, but rely on the identification of *k*-mers, short exact matching substrings of fixed length *k*, in order to compare two nucleotide sequences. For the taxonomic assignment of reads, these programs typically preprocess the reference genomes by extracting all existing *k*-mers and storing them in the index for fast lookup. Then, the *k*-mers contained in each sequencing read are searched in this index and the read is assigned to a taxon based on the matching genomes. Recent programs following this paradigm are LMAT [2], Kraken [25], and Clark [19]. For example, Kraken builds an index from all *k*-mers found in the reference genomes and assigns each *k*-mer to the least common ancestor of all species having that *k*-mer. Then, during the search, Kraken matches the *k*-mers found in the reads to this index and eventually assigns the read to the taxon with most matching *k*-mers by following a path from the root of the tree. Clark on the other hand only uses discriminative *k*-mers between sets of reference genomes belonging to a pre-defined taxonomic rank, *e.g*., genus, which are then used to classify reads to a node in the taxonomic tree at that particular rank. While this approach reduces the size of the index, it, however, prohibits the assignment of reads to higher taxonomic levels in case of ambiguity and therefore requires the user to build different indices for each rank in the taxonomic tree.

These fast methods have so far been restricted to classification at the DNA level where the fundamental requirement is a high sequence identity between reads and the reference database, so that in the minimal case at least one *k*-mer per read can be found in the database. Therefore, these methods work best for samples in which the majority of the species have been previously sequenced and their genomes are contained in the reference database and when a classification at the deepest possible level in the taxonomy is of importance. However in many samples, no reference genomes are available for a large fraction of the organisms. In such samples, a classification on the protein level is much more sensitive, because protein sequences are more conserved than the underlying DNA, and microbial and viral genomes are typically densely packed with protein-coding genes [3, 9].

Another general problem with metagenomic sequence comparison is a sampling bias in the phylogenetic distribution of available reference genomes. On the one hand, certain model organisms or pathogens, for example from human microbiomes, are primary targets for microbial research and are therefore over-represented in the genome databases. On the other hand, species that were not possible to culture in the laboratory are underrepresented, which is a further challenge for the taxonomic classification of environmental samples, especially from extreme environments. Additionally, the rate of evolution is faster for microbes and especially for viruses compared to eukaryotes due to higher replication rates. Thus, metagenomic studies continuously find novel habitats where large fractions of the sequence data remain unclassified or only show low sequence similarities to the known species [17, 23].

For these reasons, there is a need for fast metagenome classifiers, which are able to detect evolutionary distant relatives of the species having reference genomes, based on amino acid sequence comparison. Several programs exist for seed-based local alignment of protein sequences, like BlastP and BlastX[1], or the faster methods using index structures like RapSearch [26] and Diamond [4]. However, these alignment programs are generally slower than the *k*-mer based methods [15], and they report all alignments to the reference database, which need to be analyzed further for taxonomic classification.

Here we present `Kaiju`, a novel program for fast taxonomic classification based on sequence comparison to a reference database of microbial proteins. We show that our approach is able to classify more reads in real metagenomic data sets and evaluate its performance in a benchmark study, which simulates the classification of a novel genome taking the sampling bias of reference databases into account.

## Results and Discussion

Kaiju translates metagenomic sequencing reads into the six possible reading frames and searches for the best matches of amino acid sequences in a given database of annotated proteins from microbial reference genomes. The underlying sequence comparison algorithm uses the Borrows-Wheeler transform of the protein database, which enables exact string matching in time proportional to the length of the query, to achieve a high classification speed.

In *k*-mer based methods, the size of *k* governs the sensitivity and precision of the search. If *k* is chosen too large, no identical *k*-mers between read and database might be found, especially for short or erroneous reads as well as for evolutionary distant sequences. If *k* is chosen too small, more false positive matches will be found. Therefore, in order to not be restricted by a prespecified *k*-mer size, Kaiju finds maximum exact matches (MEMs) between reads and database to achieve both a high sensitivity and precision. Reads are directly assigned to species/ strain level, or in case of ambiguity, to higher level nodes in the taxonomic tree. For example, if a read contains an amino acid sequence, which is identical in two different species of the same genus, then the read will be classified to this genus. Kaiju also offers the possibility to extend matches by allowing a certain number of amino acid substitutions at the end of an exact match in a greedy heuristic approach using the BLOSUM62 substitution matrix. See Materials & Methods for a detailed description of Kaiju’s algorithm.

### Genome exclusion benchmark

Benchmarking a classifier’s accuracy can be done by simulation studies, which, knowing the ground truth about the simulated sequences, can assess the sensitivity and precision of the classification. However, the benchmark protocol needs to reflect the real obstacles in metagenomic studies, which do not only include the bias and errors of the sequencing technology, but also the microbial composition of the sample at hand. Thus, we devised a simulation benchmark, which emulates the often limited availability of reference genomes and its impact on the classification performance when faced with a novel strain or species found in the metagenomic sample. To this end, we created a reference database of 2 724 bacterial and archaeal genomes and selected the subset of genomes belonging to genera, which have at least two and most 10 genomes in the database. For each of the 882 genomes in this subset, we simulated four sets of Illumina and one set of Roche/454 sequencing reads and created a version of the reference database excluding that genome. This stripped reference (now containing 2 723 genomes) is then used to classify the simulated reads and we measure the number of classified reads, sensitivity and precision on genus as well as phylum level (see Materials & Methods). The number of genomes per genus serves as an indicator for the difficulty of the classification problem. For example, it is much harder to assign a novel genome to its genus when there is only one other genome of the same genus already available in the database. On the other hand, if there are ten genomes available in a genus, it is typically much easier to classify the reads from the excluded genome to its genus with nine remaining genomes available.

We compared the performance of Kaiju to the two *k*-mer based programs Kraken and Clark, which performed best in speed and accuracy in a recent benchmark study [15]. While Kraken uses a default length of *k* = 31, the user can chose *k* in Clark during database construction and values of *k* = 20 and *k* = 31 are recommended for highest sensitivity and highest precision respectively. Therefore we chose values of *k* = 20 and *k* = 31 in Clark in order to illustrate the influence of the choice of *k* on the classification performance. Kaiju was run in the fastest MEM mode (with minimum fragment length *m* = 11) as well as in the heuristic Greedy mode (with minimum score *s* = 65), allowing either only one (Greedy-1) or up to five (Greedy-5) amino acid substitutions during the search.

Genomes are binned into categories in the range 2 to 10 according to the total number of genomes in the genus. Sensitivity and precision are calculated as the mean across all genomes in each category for each program and the five different types of simulated reads. Figure 1 compares the genus-level sensitivity and precision and Suppl. Fig. 1 shows the mean percentage of classified reads for each genus category.

**Figure 1:**
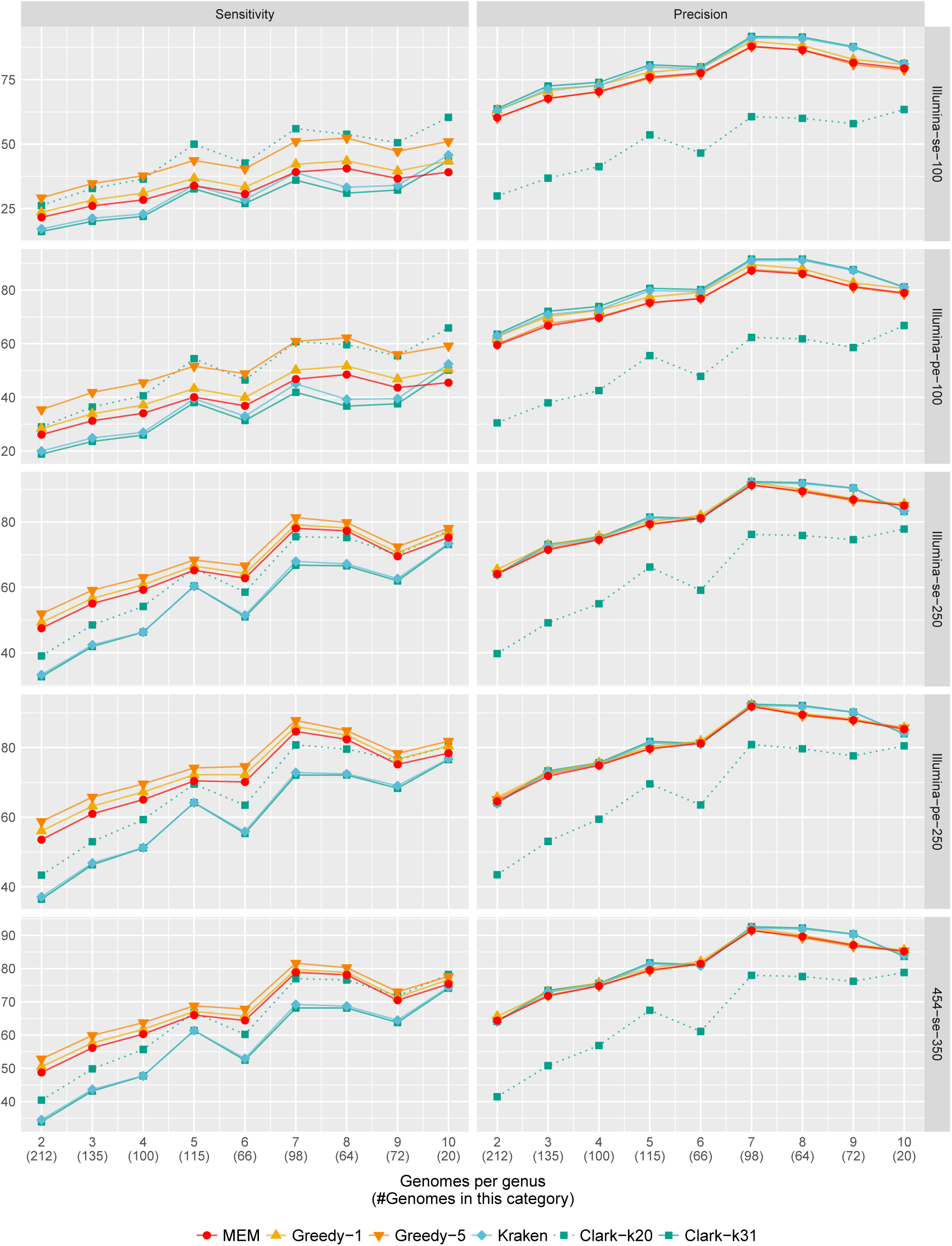
Genome exclusion benchmark: Genus-level sensitivity and precision for the five different types of simulated reads. The x-axis denotes the number of genomes in the genus and the total number of genomes in that category. For example, 212 of the measured 882 genomes belong to the 106 genera with only two available genomes, and the data points show the mean sensitivity and precision across all 212 genomes in that category. Clark with *k* = 20 is denoted by the dotted line.

As expected, all programs have the lowest percentage of classified reads and the lowest sensitivity in those genera with only few available genomes and highest sensitivity in genera with seven or more genomes. Second, the read length is a major determinant for sensitivity as there is a much higher chance of finding a matching sequence to the reference database with increasing read length. Especially Kaiju gains a further increase of sensitivity from longer reads, as the chance of an overlap to a protein-coding region additionally increases with read length. For example when looking at the Illumina single-end 100nt reads, Greedy-5 achieves the highest sensitivity of 29% of all programs in genera with only two genomes, whereas Clark-k31 has the lowest sensitivity of 16%. In contrast for Illumina paired-end 250nt reads, Greedy-5 achieves 59% sensitivity, whereas Clark-k31 only achieves 36%. With increasing number of genomes per genus, the difference between Greedy-5 and both Clark and Kraken shrinks to a few percent, as the chance of finding at least one *k*-mer per read increases with more available reference genomes. Kaiju’s MEM mode has lower sensitivity compared to Greedy modes in all cases, because it only searches for exact matches, which is especially visible in short reads.

Similarly, the precision of all programs is lowest in genera with only two genomes and increases with higher number of available genomes. However, the differences between the programs is much smaller compared to sensitivity, with Clark-k31 showing the highest precision by a small margin in most cases. When comparing Clark-k31 and Kraken-k31, Clark has consistently a little bit lower sensitivity and a bit higher precision than Kraken The difference between Clark-k20 and Clark-k31 nicely illustrates the trade-off between sensitivity and precision depending on the *k*-mer size. Generally, the loss in precision is consistently higher than the gain in sensitivity when using *k* = 20.

Suppl. Fig. 2 shows the phylum-level sensitivity and precision. At this level, the difference in sensitivity between Kaiju and Kraken is generally higher, because more reads are assigned to ranks higher than genus by Kaiju’s LCA algorithm, whereas Kraken’s weighted path algorithm usually assigns reads to the lowest possible level. Again, the increase in sensitivity with increasing read length is higher in Kaiju compared to Kraken and Clark. For example in genera with only two genomes, Greedy-5 achieves between 41% (Illumina se-100nt) and 84% (Illumina pe-250nt), whereas Clark achieves between 17% and 44%. On phylum-level, all modes of Kaiju achieve around 10% higher sensitivity than Clark and Kraken even in the highest category with genera containing 10 genomes.

**Figure 2:**
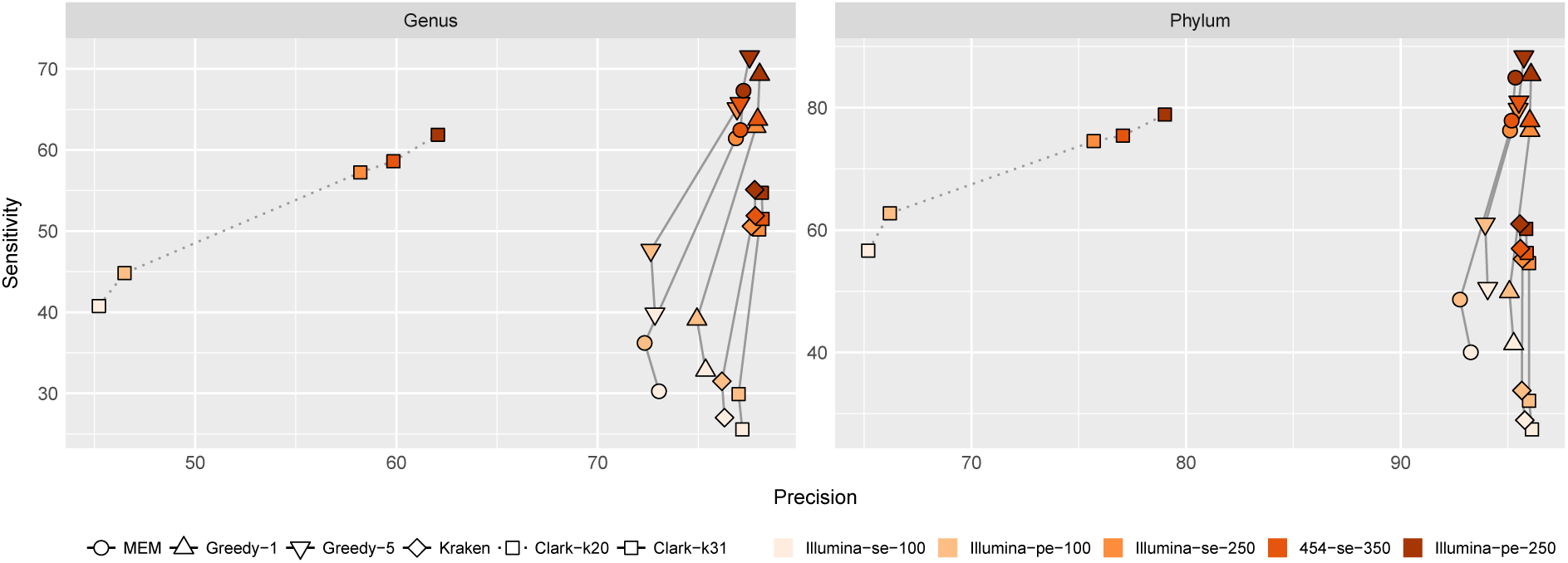
Sensitivity and precision averaged over all 882 measured genomes for the five types of reads.

Phylum-level precision is generally much higher (> 90%) for all methods and all read types compared to genus-level, because the chance of false positive matches outside the correct phylum is lower. Again, Clark-k20 consistently yields a much lower precision compared to Clark-k31 and the other programs, however it also gains more sensitivity on phylum-level classification compared to genus-level. This can be attributed to the removal of *k*-mers that are shared across genera for the genus-level classification, which, however, can be used on the phylum-level.

Figure 2 shows the mean genus-level and phylum-level sensitivity and precision across all 882 measured genomes for the five different read types. The biggest gap for sensitivity and precision between the read types occurs for all programs between both paired-end and single-end 100nt and the single-end 250nt Illumina reads. Highest sensitivity is achieved by Greedy-5, followed by Greedy-1, MEM, Kraken, and Clark in the paired-end 250nt reads. Precision is highest for Clark-k31 closely followed by Kraken both on genus-level and phylum-level. Especially in the 100nt reads, Kaiju’s precision is lower, but the gain in sensitivity remains higher than the loss in precision. For the 250nt reads and longer, Kaiju’s precision is marginally lower than Kraken and Clark-k31, while sensitivity is much higher. Clark-k20 shows lowest precision in all read types compared to the other programs both on genus and phylum levels.

In this analysis, we used cutoff values of minimum required match length *m* = 11 in Kaiju’s MEM mode and minimum required match score *s* = 65 in the Greedy modes. Suppl. Fig. 3 shows the tradeoff between sensitivity and precision of the classification depending on the choice of *m* or *s*. Similar to the choice of *k* in Clark, the sensitivity is highest and precision is lowest for small cutoff values. Increasing the cutoffs results in lower sensitivity but higher precision. However, the increase in sensitivity between *m* =11 and *m* = 12 is higher than the loss in precision in all datasets both on genus as well as phylum-level. Similarly in the Greedy modes, *s* = 65 also yields higher gain in sensitivity than loss in precision.

### Real metagenomes

In order to assess how many reads can actually be classified in real metagenomic datasets, we arbitrarily selected ten datasets from different microbiomes, which were sequenced using various different HTS instruments. The two datasets from human saliva and vagina samples were already used in [19]. The other eight samples are derived from human and cat gut, a freshwater lake, the Amazon river plume and Baltic sea, xeric desert soil and from two bioreactors, which were inoculated with microbes from Wadden Sea sediment and compost environments. Suppl. Tab. 1 lists metadata and accession numbers for the datasets. The same database comprising 2 724 genomes from our exclusion benchmark serves as a reference database. We classified the ten datasets using Kraken (*k* = 31) and Kaiju in MEM and Greedy-5 modes with more conservative cutoff values of *m* =12 and *s* = 70 respectively, which showed on average a similar precision as Kraken across the five types of reads in our exclusion benchmark, see Suppl. Fig. 3. We also mapped the four human and cat samples to their respective host genomes using BWA [14], and the percentage of mapped reads was at most 2%.

Figure 3 shows the percentage of classified reads from each dataset for MEM, Greedy-5, and Kraken, as well as the overlap and combined percentage of Greedy-5 and Kraken. Generally, Kaiju’s MEM mode classifies between 13.1% (Human Vagina) and 48.8% (Bioreactor Sediment) more reads than Kraken, which is further increased to 17.8% and 56.6% in Kaiju’s Greedy-5 mode. The percentages of reads that are classified by Kraken, but unclassified by Greedy-5 range between 0.3% (Desert Soil and Lake) and 4.4% (Human Gut). Across all datasets, the number of reads that were classified by both Greedy-5 and Kraken (overlap) varies between 2.8% (Lake) and 42.4% (Human Vagina). By merging the results from Greedy-5 and Kraken, between 24.7% (Desert Soil) and 73.1% (Bioreactor Sediment) of the total reads can be classified.

**Figure 3:**
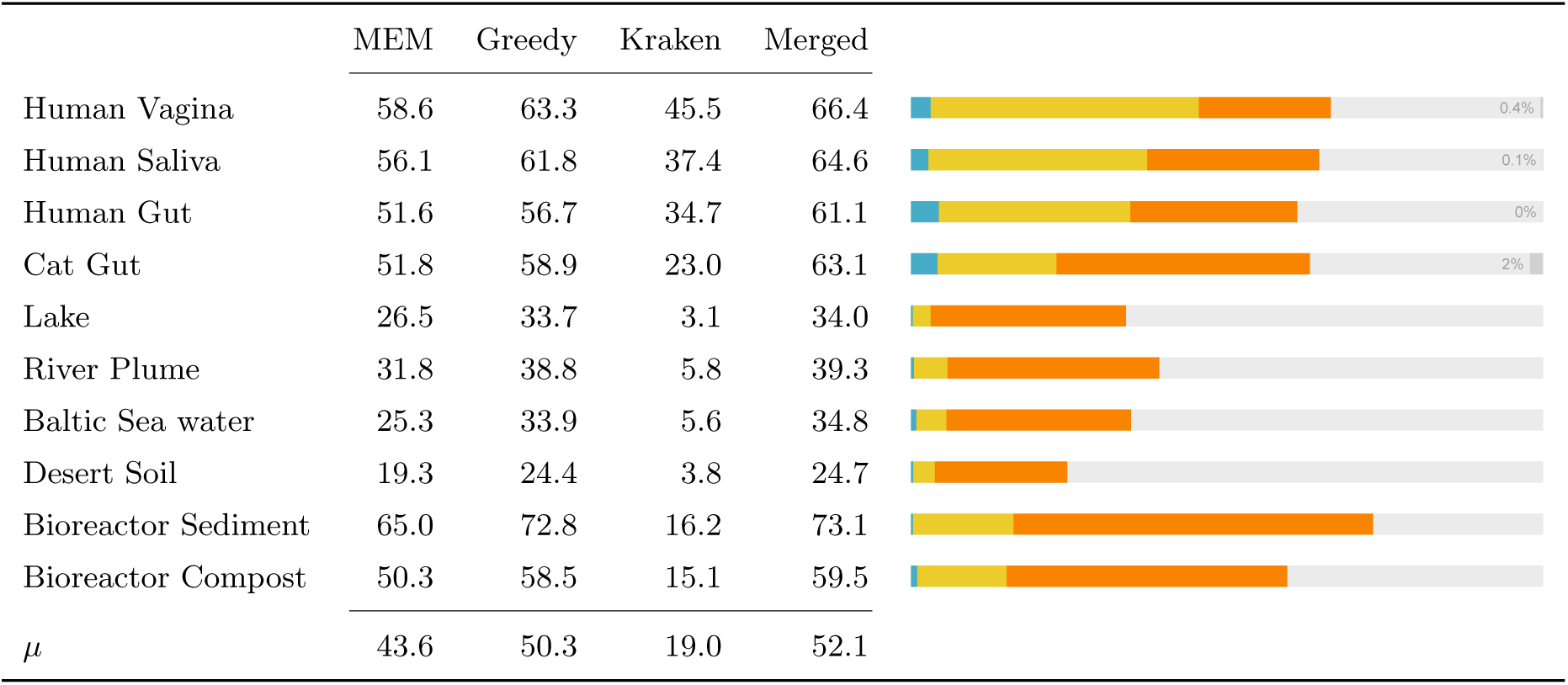
Percent of classified reads in the ten real metagenomes for Kaiju MEM (*m* = 12) and Greedy-5 (*s* = 70) as well as Kraken (*k* = 31). The *Merged* column shows the percentage of reads that are classified by at least one of Greedy-5 or Kraken. The Venn-Bar-diagram visualizes the percentage of reads that are classified either only by Kraken (blue), Greedy-5 (orange), or both (yellow). Gray bars in the human/cat samples denote the percentage of reads mapped to the respective host genomes.

As expected, the environmental samples, especially from the extreme xeric desert, but also the aquatic microbiomes, pose the biggest challenges for taxonomic assignment. In those samples Kaiju’s protein-level comparison with substitutions allows for a more sensitive sequence comparison resulting in more classified reads. However, even in the human microbiomes Kaiju’s protein-level classification adds more than 20% additional classified reads to Kraken’s result.

We also run each dataset through Clark (*k* = 31) with its phylum-level database and it classified fewer reads than Kraken in all cases (data not shown). In principle, if only a small fraction of reads were classified, classification could be done using a smaller *k*-mer size in Kraken and Clark or smaller cutoff values in Kaiju in order to increase the number of classified reads. The trade-off, however, would be a decreased precision as shown in our benchmark and also discussed in [19].

### HiSeq and MiSeq mock communities

Additionally to the real metagenomes, we also measured Kaiju’s and Kraken’s performance using the same metrics and reference database on the HiSeq and MiSeq mock community datasets that were used in [25, 19]. They comprise 10k real sequencing reads from 10 bacterial strains with mean read length of 92nt (HiSeq) and 156nt (MiSeq). All strains belong to genera that are associated with human microbiomes or human pathogens and have typically many reference genomes available. Suppl. Tab. 2 shows sensitivity and precision on both genus and phylum-level of Kaiju in Greedy-5 mode and Kraken (*k* = 31) using the same reference database as above. In the HiSeq dataset, Kaiju has 73.3% sensitivity (Kraken: 78.0%) and 94.4% precision (Kraken: 99.2%) on genus level, and 78.1% sensitivity (Kraken: 78.8%) and 98.3% precision (Kraken: 99.7%) on phylum level. Because the short reads can only yield short amino acid fragments, which are more likely found across genera, many reads are assigned to higher ranks resulting in a lower genus-level sensitivity. Additionally, the short read length results in generally lower overlap with protein-coding regions and therefore Kraken yields a higher sensitivity, because it can classify those. In the MiSeq dataset, the difference between both programs on genus level is similar, whereas Greedy-5 yields 8% higher sensitivity and 1% higher precision on phylum-level compared to Kraken.

### Runtime and Memory

The dataset for the runtime benchmark contained 27.24m reads comprised of 10k randomly sampled reads from each of the 2 724 genomes in our accuracy benchmark, which served again as the reference database. For the five different types of reads, the classification speed of Clark and Kraken using *k* = 31 and of Kaiju’s modes MEM, Greedy-1, and Greedy-5 was measured using 25 parallel threads (see Materials & Methods for specification of the hardware).

Figure 4 shows the number of processed reads per second (rps). The classification of the short singleend Illumina 100nt reads is the fastest in all programs (Kaiju MEM: 166k rps, Kraken: 165k rps, Clark: 93k rps), whereas classification of the Illumina paired-end 250nt (MEM: 94k rps, Kraken: 24k rps, Clark: 19k rps) takes the longest time. In the long reads, Kaiju can benefit from search space pruning by finding long MEMs first, whereas Kraken and Clark have to analyze more *k*-mers compared to the shorter reads. Naturally, Kaiju’s MEM mode is much faster than the Greedy modes, which extend the search space and also need to calculate the scores for each match. Depending on the read type, Greedy-5 classifies between 15k and 41k rps. Greedy-1 with only one allowed mismatch is faster than Greedy-5 and can classify between 34k and 69k rps. Interestingly, Kaiju’s Greedy mode is faster in longer reads compared to the 100nt reads. This is due to the pruning of the search space by discarding query sequences that cannot achieve higher scores than the best scoring match, which is usually found earlier in longer reads (see Materials & Methods). While Kaiju MEM is the fastest program in most cases, especially for the short reads, Kaiju’s Greedy-5 generally takes the longest time, nicely demonstrating the trade-off between speed and sensitivity.

**Figure 4:**
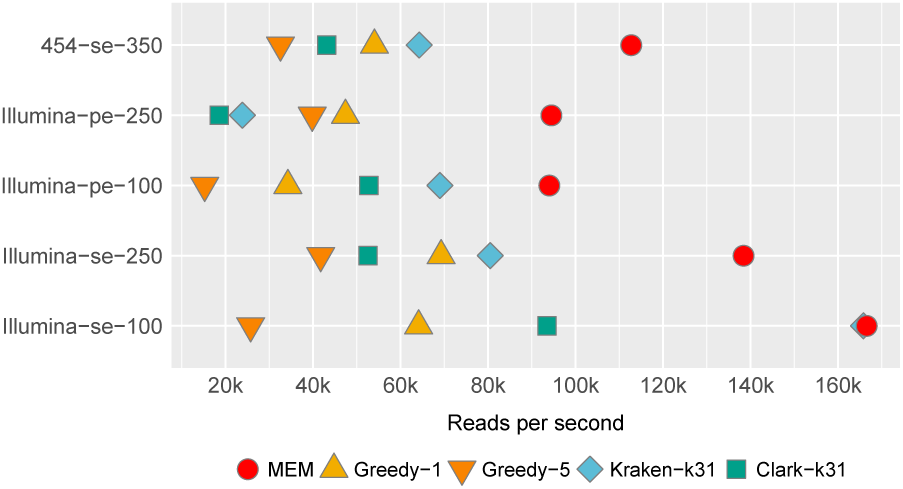
Classification speed in processed reads per second using 25 threads for the five read types.

The measured peak memory consumption during the classification is 5.6 GB for Kaiju, 72 GB for Kraken and between 65 and 78 GB for Clark depending on the read length.

The construction of Kaiju’s database index from the protein sequences takes 8 minutes with peak memory usage of 24 GB using 25 threads (Kraken: 1h26m/165 GB, Clark: 3h57m/152 GB). The memory usage is largely due to the parallel sorting of the suffix array and can be reduced by using fewer parallel threads, *e.g*., only 6.6 GB are needed with 5 threads. Kaiju’s final index size on disk is 4.8 GB (Kraken: 73 GB, Clark: 39 GB).

## Conclusion

When performing sequence comparison, as in the case of taxonomic assignment using a reference database, there is the obvious trade-off between an algorithm’s speed and accuracy. While the traditional local alignment would return optimal alignments, its slow runtime prohibits its use on large HTS datasets. On the other hand, while *k*-mer based methods are very fast, they often lack sensitivity and a big fraction of the metagenomic reads might remain unclassified, as can be seen from Figure 3. Kaiju therefore uses maximum exact matches (with optional substitutions) on the protein-level instead nucleotide-level in order to increase sensitivity of the classification while maintaining a high precision. By using the Borrows-Wheeler transform as an index for the reference protein database, Kaiju is still fast enough for classifying up to millions of reads per minute, depending on the read length and the number of allowed mismatches. Additionally, Kaiju’s memory footprint is small enough (below 6 GB) for running the program on a standard PC/laptop.

The aim of using protein-level sequence comparison is to improve the classification of metagenomes comprising species that are evolutionary distant to the species in the reference or belong to genera that have only few reference genomes available. Therefore, we focused on those genera with ten or less genomes in our genome exclusion benchmark, because the classification problem becomes easier once many reference genomes are available and can be mostly accomplished by nucleotide-level sequence comparison, which is also better suited for strain typing because of the finer resolution regarding SNPs. Our benchmark on 882 genomes and five types of simulated reads shows that Kaiju consistently achieves a much higher sensitivity with only little loss of precision compared to Kraken and Clark, which use fixed-length *k*-mers. The difference was especially visible in genera with only few available genomes.

The obvious disadvantage of protein-level sequence classification is the inability to classify reads originating from non-protein-coding genomic regions. Especially when the genomes of the sequenced microbial strains are also contained in the reference database, Kaiju would be less sensitive than nucleotide-level classifiers, which can assess the entire genome, as seen in the HiSeq and MiSeq datasets. However, due to the high density of protein-coding genes in microbial genomes, the probability of overlap between individual sequencing reads and protein-coding genes increases substantially with increasing read lengths. Furthermore, when using paired-end sequencing, the chance of one mate overlapping with a protein-coding gene is higher than for single-end sequencing, which was also shown in our benchmark where longer and paired-end reads had higher sensitivity compared to shorter single-end reads.

In our set of ten randomly selected real metagenomic datasets, Kaiju consistently classifies on average twice as many reads as Kraken. The highest differences are observed in samples from nonhuman microbiomes, showing that especially the classification of environmental samples with high evolutionary distances to reference genomes can gain from Kaiju’s more sensitive sequence comparison. By combining Kaiju’s and Kraken’s output, between 24% and 73% of reads can be classified across the various samples.

Principally, Kaiju’s algorithm is not limited to assigning reads to taxa, but can also be used for fast searching in arbitrary protein databases, for example when querying novel bacterial genomes against a database of resistance genes. The certainly expected increase of reference database volumes in the coming years can easily be handled by Kaiju, due to the usage of a sparse suffix array and FM-index, resulting in a small memory footprint.

## Materials and Methods

### Metagenome classifier

Kaiju classifies individual metagenomic reads using a reference database comprising the protein-coding genes of a set of microbial genomes. We employ a search strategy, which finds maximal exact matching substrings (MEMs) between query and database using a modified version of the backwards search algorithm in the Borrows-Wheeler Transform [6, 8]. The Borrows-Wheeler Transform (BWT) [5] is a text transformation that converts the reference sequence database into an easily searchable representation, which allows for exact string matching between a query sequence and the database in time proportional to the length of the query. Whereas in the context of read mapping, MEMs have been used as a fast method for identifying seeds of mapping regions in the reference genome, for example in [16, 13], we use MEMs to quickly find those sequences in the reference database, which share the longest possible subsequence with the query. Lookups in the BWT are performed using a checkpointed variation of the FM-index [6], which allows for decreasing the otherwise large memory requirement due to the size of the amino acid alphabet. The initial suffix array used for calculating the FM-index is saved as a sparse suffix array, which further reduces the size of the database index with only little impact on runtime, because the suffix array is only needed to extract the name of the database sequence once the best match for a read is found. Upon index creation, the distance between suffix array check-points can be chosen by the user, allowing to trade-off memory usage and search speed.

First, Kaiju translates each read into the six possible reading frames, which are then split at stop codons into amino acid fragments. These fragments are sorted by length, and, beginning with the longest fragment, queried against the reference database using the backwards search in the BWT (Figure 5). Given a query fragment of length *n* and the minimum required match length *m*, the backwards search is started from all positions between *n* and *n* – *m* in the query and the longest MEMs are retained. If one or more matches of length *l* > *m* are found, *m* is set to *l* and the next fragment is queried against the database if its length is at least *l*, otherwise the search stops. Once the search is finished and one or more matches of maximum length are found, the taxon identifiers from the corresponding database sequences are retrieved from the suffix array and printed to the output. If matches are found in multiple taxa, Kaiju determines their least common ancestor and outputs its taxon identifier. Thus, each read is always classified to the lowest possible taxonomic level given the ambiguity of the search result.

**Figure 5:**
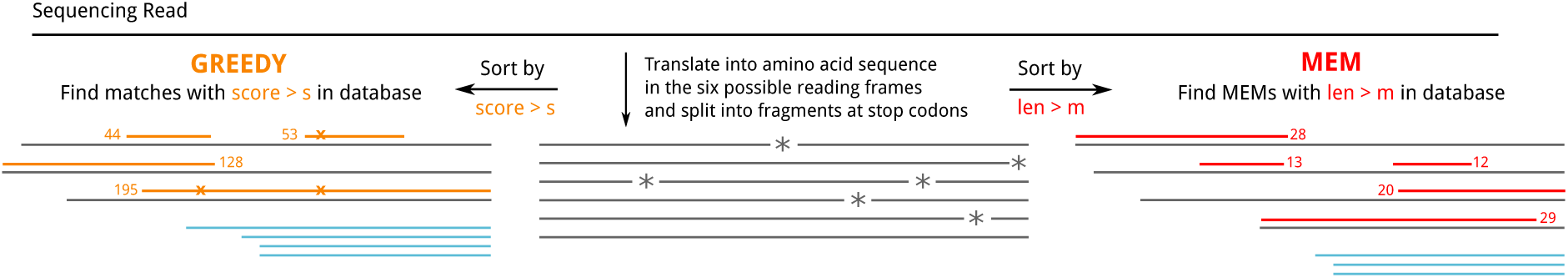
Kaiju’s algorithm for maximum exact matching (right side) and greedy matching (left side). Grey lines denote the translated amino acid fragments that are searched in the database, whereas red and orange lines indicate matches in MEM and Greedy mode. Blue lines indicate fragments that are not evaluated because their length or maximum obtainable score is lower than the length or score of the best match from previously searched fragments.

The minimum required MEM length *m* is the major parameter for trading sensitivity versus precision (with little impact on runtime). If the error rate *e* of the sequencing reads is known and the evolutionary distance between reference genome and sequenced genome is negligible, *m* can be estimated from *e* and the read length [16]. However, in metagenomics, the evolutionary distance, which adds variation on top of sequencing errors, is not known a priori. At least, one can estimate the false positive rate by counting random matches. To this end, we created a shuffled version of the microbial subset of NCBI’s *NR* protein database, using uShuffle [11] with a window length of 100 amino acids, and searched for MEMs between simulated metagenomic reads and the shuffled database. Suppl. Fig. 5 shows the cumulative sum of random matches sorted by the length of the match, and one can observe that ~95% have length ≤11. When classifying simu-
lated reads against the original database, more than 75% of wrong classifications and only ~2% of correct classifications have length ≤ 11. We therefore set *m* = 11 as the default minimum match length in Kaiju.

Searching for MEMs is the fastest possible search strategy, but its sensitivity decreases with increasing evolutionary distance between query and target, where more and more amino acid substitutions occur and exact matches become shorter. Therefore, allowing for substitutions during the backwards search can bridge mismatches and extend the match at the cost of an exponential increase of runtime depending on the number of allowed mismatched positions. Because of the rapid expansion of the search space, especially with the 20 letter amino acid alphabet, one could employ a greedy heuristic, in which substitutions are only introduced at the end of a match instead of all positions in the query sequence. Therefore, we also implemented a *Greedy* search mode in Kaiju, which first locates all MEMs of a minimum seed length (default 7) and then extends them by allowing substitutions at the left ends of each seed match. From there, the backwards search continues until the next mismatch occurs. Eventually the search stops once the left end of the query is reached or if the maximum allowed number of substitutions has been reached.

Since amino acid substitutions in homologous sequences are non-uniform, a further speed-up can be gained by prioritizing the most likely substitutions at each position. By using an amino acid substitution model, a total score for each match can be calculated, as in standard sequence alignment, which is then used to rank multiple matches and select the taxon from the database for classification. Therefore, after the translation of a read into a set of amino acid fragments, we rank the fragments by their BLOSUM62 score and start the database search with the highest scoring fragment. For each substituted amino acid the modified fragment is placed back into the search list according to its new (now lower) score. Once a match is found, which has a higher score than all remaining fragments in the search list and a score above the minimum score threshold *s*, the algorithm stops and this highest-scoring match is used for classifying the read. Again, the minimum required score *s* necessary for avoiding random matches can be estimated by using a shuffled database and we use *s* = 65 as default value for Kaiju’s Greedy mode.

Kaiju is implemented as a command-line program in C/C++. It reads input files in either FASTA or FASTQ format containing the (single-end or paired-end) reads and outputs one line for each read (or read pair), containing the read name and the NCBI taxon identifier of the assigned taxon as well as the length/score of the match. Optionally, Kaiju can also produce a summary file with the number of reads assigned per taxon, which can be loaded into Krona [18] for interactive visualization. We also include a utility program, which can merge the classification results from different runs or programs, *e.g.* for merging Kaiju and Kraken results.

### Performance evaluation

The primary goal of Kaiju’s protein-level classification is to improve classification of those parts of a metagenome that are only distantly related to the known sequences or belong to a branch of the phylogeny that is underrepresented in the reference database. We therefore devised a benchmark study, which addresses this problem by simulating the classification of metagenomic reads from a novel strain or species, which is not contained in the reference database.

For our benchmark dataset, we downloaded a snapshot of all complete bacterial and archaeal genomes from the NCBI FTP server (date: 2014–12–16). Only those genomes were retained that are assigned to a species belonging to a genus and have a full chromosome with annotated proteins, resulting in a total of 2 724 genomes belonging to 692 distinct genera. Suppl. Fig. 4 shows the distribution of genomes to genera, illustrating the large variance in the number of sequenced genomes for each genus. For example, the genus *Streptococcus* contains 121 genomes, whereas 405 genera have only one available genome, 106 genera have two available genomes, and so on. The distribution clearly illustrates a sampling bias and the sparseness across large parts of the phylogeny.

From the total of 2 724 genomes, we extracted those genera, which have at least 2 and at most 10 genomes assigned. This resulted in a list of 242 genera comprising 882 genomes, for which we measure the classification performance individually. For each of the 882 genomes we simulated five sets of HTS reads and created a reference database not containing this genome, which is then used to classify the simulated reads. Reads were simulated from the whole genome (including plasmids) using ART [10]. The four sets of Illumina reads contain 50k reads of length either 100nt or 250nt, both in single-end and paired-end mode. Another set of 50k Roche/454 reads with minimum length of 50nt and mean length of 350nt was also simulated using ART.

To evaluate classification accuracy, we measure the number of classified reads as well as sensitivity and precision on genus and phylum levels. Sensitivity is calculated as number of reads assigned to the correct genus/phylum divided by the total number of reads in the input. Precision is calculated as the number of reads assigned to the correct genus/phylum divided by the number of classified reads minus the number of reads that were classified correctly to a rank above genus level. The same measurements were used in [25]. Kraken (ver 0.10.4b) and Clark (ver 1.1.3) were run in their default modes using *k* = 31 for highest precision, and Clark was also run using *k* = 20. Kaiju was run in MEM mode using *m* = 11 … 14 and in Greedy-1 and Greedy-5 modes (allowing only 1 or up to 5 substitutions) using *s* = 55 … 80.

Speed measurements were run on an HP Apollo 6000 System ProLiant XL230a Gen9 Server, which has two 64-bit Intel Xeon E5–2683 2GHz CPUs (14 cores each), 128GB DDR4 memory and a 500GB 7200rpm SATA disk (HP 614829–002). Kraken and Clark were run in default modes with *k* = 31 and Kaiju was run in MEM (*m* = 12) as well as Greedy-1 and Greedy-5 (*s* = 65) modes. Performance was measured in processed reads (or read pairs) per second (rps) using 25 parallel threads. While Kaiju and Clark need to preload to index into memory before the classification starts, Kraken can either preload the index or only load necessary segments during the classification. We therefore measured Kraken’s speed using both options, and it turned out that Kraken runs faster without preloading on our hardware. We therefore report its performance without preloading. For each of the five types of simulated reads from our exclusion benchmark, we created a dataset comprising 10k reads from each genome in the reference database, resulting in 27.24m reads for each read type. Each combination of program and read type was measured four times to reduce impact of caching and I/O fluctuations and the fastest run of the replicates is reported.

## Acknowledgments

The research leading to these results has received funding from the European Union 7th Framework Programme FP7/2007–2013 under grant agreement nr. 265933.

